# The digital sphinx: Can a worm brain control a fly body?

**DOI:** 10.64898/2026.03.20.713233

**Authors:** Bingni W. Brunton, Elliott T.T. Abe, Lawrence Jianqiao Hu, John C. Tuthill

**Affiliations:** Dept of Biology, University of Washington, Seattle, WA, USA; Graduate Program in Neuroscience, University of Washington, Seattle, WA, USA; Dept of Neurobiology & Biophysics, University of Washington, Seattle, WA, USA

## Abstract

Animal intelligence is not purely a product of abstract computation in the brain, but emerges from dynamic interactions between the nervous system and the body. New connectome datasets and musculoskeletal models now enable integrated, closed-loop simulations of the neural and biomechanical systems of the fruit fly *Drosophila*, an ideal model organism to investigate embodied intelligence. However, many biological parameters of the nervous system and the body, as well as how they interface, remain unknown. To fill such gaps, researchers are turning to deep reinforcement learning (DRL), a data-driven optimization framework, to create virtual animals that imitate the behavior of real animals. Here, we provide a cautionary tale about the interpretation of such models. We constructed a virtual chimera of two phylogenetically distant species: a connectome of the *C. elegans* nematode worm and a biomechanical model of the fly body. The worm connectome receives sensory information from the fly body, and an artificial neural network is trained with DRL to map worm motor neuron activations to the fly’s leg actuators. The resulting digital sphinx produces highly realistic fly walking—yet it is biologically meaningless. This exercise teaches us nothing about either animal and exposes a core peril of connectome-body models: behavioral fidelity is achievable without biological fidelity, making such models easy to overinterpret. Done carefully, virtual animals can be powerful partners to biological experiments, but only if their components and interfaces are grounded in biology.

---

In the last few years, researchers in neuroscience and AI have been working towards developing computational models of brains capable of controlling realistic animal bodies in virtual environments (e.g., [1, 2, 3, 4, 5, 6]). These virtual animal models have the potential to enable *in silico* experiments at scale, generating new theories and hypotheses about embodied intelligence to be tested in the lab. They could also serve as community repositories of accumulated knowledge about the nervous system and the body. Motivated by the potential utility of such models, the field of virtual animals is growing rapidly, including both academic and industry research efforts. Some have referred to these models as brain “emulations” or “uploads,” attracting substantial interest from scientists and the public^1^. But such models of virtual animals require cautious interpretation.

Among animals that walk, the integration of brain wiring and body models is perhaps closest to fruition in *Drosophila*, due to the recent completion of multiple complete wiring diagrams (known as connectomes) of the fly nervous system. The fly is the only animal with legs for which nearly comprehensive connectomes of its brain and nerve cord exist [7, 8]. There is also a wealth of knowledge about the neural basis of many fly behaviors, along with established tools for experimentally recording and manipulating genetically identified cell types in the fly nervous system. Two open-source biomechanical models of the whole fly body [9, 3] have been implemented in the physics engine MuJoCo [10]. Despite this progress, closed-loop integration of biomechanical and neural models remains far from straightforward. Key parameters governing how they interface remain incompletely characterized, including how motor neuron commands result in forces exerted by the body and how sensory neurons detect and transduce internal and external stimuli.

Where interfaces between brains and body models are missing or only partially characterized, one approach is to train an artificial neural network (ANN) to approximate these interfaces with deep reinforcement learning (DRL). In building virtual animal models, a motor policy is commonly learned by DRL so that the integrated, closed-loop virtual body successfully mimics the detailed kinematics of real animal behavior. The implicit logic is, if it looks and acts like a cat, it is a decent model of a cat.

Additional realism is added when the motor policy network is constrained by a connectome dataset. However, many biophysical parameters for individual neurons and synapses remain unmeasured. To approximate these, researchers have used neurophysiological recordings, optogenetic perturbation experiments, and other biological observations as additional constraints [11, 12, 13, 14]. These approaches have demonstrated success at predicting some key aspects of neural dynamics, such as sensory receptive fields or motor rhythms, which will be essential for developing meaningful closed-loop models based on connectome datasets.

## Methods

Here we develop a virtual chimera with the brain of a worm and the body of a fly. We used the adult hermaphrodite *C. elegans* nematode connectome dataset [15, 16, 5], including the identities of its 302 neurons and their synapses (**Fig. 1A**). We created a graded activation model of the network, consistent with the fact that most *C. elegans* neurons do not fire action potentials. Following the model in [5], synaptic weights were based on the number of synapses from the connectome, signed by the neurotransmitter identity of each presynaptic neuron. To simulate the physics of fly walking, we used a MuJoCo biomechanical model of the fly body [3]. The fly body model had 42 actuators for its 6 legs that were driven by torque commands; proprioceptive feedback from 148 sensors reported leg and body angles, along with their derivatives.

**Figure 1:**
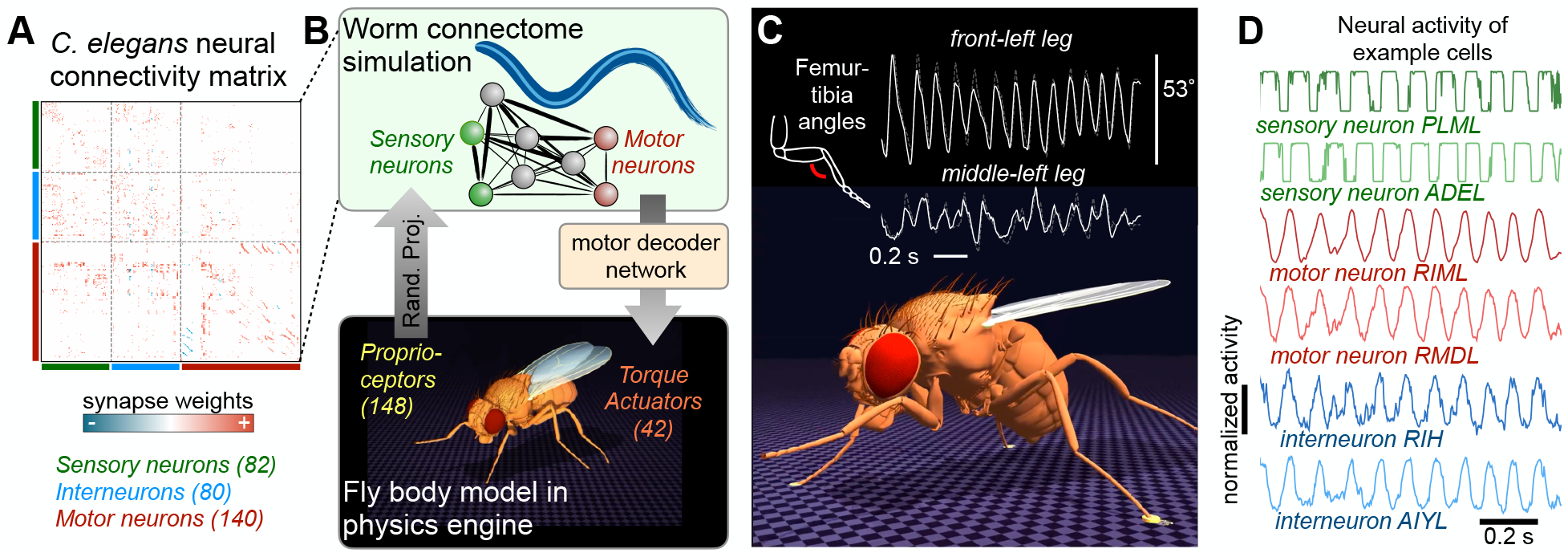
The digital sphinx model had the brain of a worm and the body of a fly, and it is able to reproduce physically realistic walking with DRL training (see walking animation in **Video 1**). **A**, Connectivity matrix from the adult hermaphrodite *C. elegans* connectome. **B**, A schematic of how the connectome simulation interfaces with the fly body model with proprioceptors and actuators. **C**, Example traces of leg joint kinematics produced by the model (white) trained to imitate target trajectories from real flies (gray dashed). **D**, Example traces of neurons in the simulated connectome model. The traces were highly cyclic because the connectome was driven by proprioceptive sensory signals in closed loop during cyclic stepping. The identities of the neurons are not biologically meaningful in this model.

To interface brain and body, we routed proprioceptive signals (normalized and filtered through a tanh nonlinearity) from the fly body to the worm’s 82 sensory neurons by random projection, without training any parameters at the sensory interface (**Fig. 1B**). We did not discriminate among different types of sensory neurons (e.g., proprioceptive, chemosensory, tactile, etc.). To actuate the body model, we then used DRL to train a variational encoder-decoder network with a 16-dimensional latent space to imitate the 3D joint kinematic trajectories of real walking flies [17, 18]. This motor decoder maps worm motor neuron activity into torque commands for the fly’s legs. The policy network was trained in closed loop using PPO as implemented by MIMIC-MJX [19].

## Results

The digital sphinx model, with a worm brain and a fly body, produced highly realistic walking, including joint angle trajectories and leg coordination patterns (**Fig. 1C, Video 1**). Yet the model is implausible and scientifically meaningless. Worms move by wiggling; flies walk with articulated legs. The fly brain is 500X larger than the nematode’s, and fly legs alone are controlled by more motor neurons than exist in the entire worm nervous system. The model produced neural activity (**Fig. 1D**), but it is uninterpretable. The model also fails to qualify as brain “emulation,” which would require that the model preserves the biological meaning of its components. The fact that a worm connectome can be trained to control fly walking reveals nothing about either worms or flies.

## Discussion

These results carry two lessons. First, users and readers beware—DRL is a capable optimization algorithm. The motion imitation approach is able to fit expressive ANNs to complex target data, even when the imposed constraints are incomplete or outright wrong. Second, even when brain and body models are individually accurate, a biologically unrealistic interface between them renders the whole model meaningless. In our digital sphinx, the worm connectome functions simply as a recurrent neural network (RNN) with rich enough dynamics to support learning of realistic locomotor trajectories. Its role in the movement policy could be fulfilled equally well by a randomly connected RNN, akin to reservoir computing [20], since all the learning happens in the black-box ANN motor decoder. This exercise is an example of how a functioning interface is not necessarily a meaningful one. Here, we did not train synaptic weights or cellular parameters of the connectome simulation; training them would only deepen this problem.

On a positive note, we are optimistic that a virtual animal that captures the coordinated function of neural and biomechanical systems can indeed be achieved, but only through careful design of its modules and interfaces. First, neuromechanical models must be well grounded in biological knowledge. Most critically, the inputs and outputs of the connectome model must be accurately connected to the muscles and sensors in the body model. This mapping has been partially achieved by combining the connectome datasets with other imaging modalities, but much more work is needed to accurately identify the thousands of sensory and motor neurons in the fly connectomes. The body models also lack biological realism in their musculature, sensory neurons, and skeletal mechanics; ongoing efforts are gradually improving these body models.

Second, neuromechanical models should be developed in close collaboration with biological experiments. Models can be powerful even without perfectly realistic components, provided they generate testable predictions about feasible experiments. Looking further ahead, swapping brain and body models of related species may one day yield real insights into how their brains and bodies diverged through evolution. However, far more model development and experimental validation is needed before we can learn anything from such a digital sphinx.

## Acknowledgements

We thank Eiman Azim, Carl Bergstrom, Astra Bryant, Michael Dickinson, and Talmo Pereira for comments on the manuscript. We thank Sarah Pugliese for assistance with parsing the *C. elegans* connectome. This work was funded by NIH grant U01NS136507 to B.W.B., R01NS14543 to J.C.T. and B.W.B., and the Richard & Joan Komen University Chair to B.W.B. J.C.T is a New York Stem Cell Foundation–Robertson Investigator.

## Author Contributions

B.W.B. and E.T.T.A. conceived of the study and designed the analyses. E.T.T.A. and J.L.H. developed and implemented the model. B.W.B. and J.C.T. wrote the manuscript.

## Data and Code Availability

All code to run the simulations, results from computational experiments, and scripts to reproduce the analyses are available on GitHub: https://github.com/Brunton-Lab/DigitalSphinx2026.

## Supplementary Video

**Video 1:** A worm brain controls a fly body and produced realistic fly walking kinematics and coordination patterns. The model also produced neural activity of cells in the worm connectome, shown here overlaid on their locations in a worm body, with a few example traces. The identities of cells in this model are meaningless and their neural activity is not interpretable.

## Footnotes

1 See https://x.com/alexwg/status/2030217301929132323

## References

[1] J. Merel, D. Aldarondo, J. Marshall, Y. Tassa, G. Wayne, and B. Ölveczky, “Deep neuroethology of a virtual rodent,” Nov. 2019. arXiv:1911.09451.

[2] D. Aldarondo, J. Merel, J. D. Marshall, L. Hasenclever, U. Klibaite, A. Gellis, Y. Tassa, G. Wayne, M. Botvinick, and B. P. Ölveczky, “A virtual rodent predicts the structure of neural activity across behaviours,” Nature, vol. 632, pp. 594–602, Aug. 2024.

[3] R. Vaxenburg, I. Siwanowicz, J. Merel, A. A. Robie, C. Morrow, G. Novati, Z. Stefanidi, G.-J. Both, G. M. Card, M. B. Reiser, M. M. Botvinick, K. M. Branson, Y. Tassa, and S. C. Turaga, “Whole-body physics simulation of fruit fly locomotion,” Nature, pp. 1–9, Apr. 2025.

[4] S. Wang-Chen, V. A. Stimpfling, T. K. C. Lam, P. G. Özdil, L. Genoud, F. Hurtak, and P. Ramdya, “Neuromechfly v2: simulating embodied sensorimotor control in adult Drosophila,” Nature Methods, vol. 21, no. 12, pp. 2353–2362, 2024.

[5] M. Zhao, N. Wang, X. Jiang, X. Ma, H. Ma, G. He, K. Du, L. Ma, and T. Huang, “An integrative data-driven model simulating C. elegans brain, body and environment interactions,” Nature Computational Science, vol. 4, no. 12, pp. 978–990, 2024.

[6] X. Liu, M. D. Loring, L. Zunino, K. E. Fouke, F. A. Longchamp, A. Bernardino, A. J. Ijspeert, and E. A. Naumann, “Artificial embodied circuits uncover neural architectures of vertebrate visuomotor behaviors,” Science Robotics, vol. 10, no. 107, p. eadv4408, 2025.

[7] A. S. Bates, J. S. Phelps, M. Kim, H. H. Yang, A. Matsliah, Z. Ajabi, E. Perlman, K. M. Delgado, M. A. M. Osman, C. K. Salmon, J. Gager, B. Silverman, S. Renauld, M. F. Collie, J. Fan, D. A. Pacheco, Y. Zhao, J. Patel, W. Zhang, L. Serratosa Capdevilla, R. J. Roberts, E. J. Munnelly, N. Griggs, H. Langley, B. Moya-Llamas, R. T. Maloney, S.-c. Yu, A. R. Sterling, M. Sorek, K. Kruk, N. Serafetinidis, S. Dhawan, T. Stürner, F. Klemm, P. Brooks, E. Lesser, J. M. Jones, S. E. Pierce-Lundgren, S.-Y. Lee, Y. Luo, A. P. Cook, T. H. McKim, E. C. Kophs, T. Falt, A. M. Negrón Morales, A. Burke, J. Hebditch, K. P. Willie, R. Willie, S. Popovych, N. Kemnitz, D. Ih, K. Lee, R. Lu, A. Halageri, J. A. Bae, B. Jourdan, G. Schwartzman, D. D. Demarest, E. Behnke, D. Bland, A. Kristiansen, J. Skelton, T. Stocks, D. Garner, F. Salman, K. C. Daly, A. Hernandez, S. Kumar, T. B.-F. Consortium, S. Dorkenwald, F. Collman, M. P. Suver, L. M. Fenk, M. J. Pankratz, G. S. Jefferis, K. Eichler, A. M. Seeds, S. Hampel, S. Agrawal, M. Zandawala, T. Macrina, D.-Y. Adjavon, J. Funke, J. C. Tuthill, A. Azevedo, H. S. Seung, B. L. de Bivort, M. Murthy, J. Drugowitsch, R. I. Wilson, and W.-C. A. Lee, “Distributed control circuits across a brain-and-cord connectome,” bioRxiv, 2025.

[8] S. Berg, I. R. Beckett, M. Costa, P. Schlegel, M. Januszewski, E. C. Marin, A. Nern, S. Preibisch, W. Qiu, S.-y. Takemura, A. M. Fragniere, A. S. Champion, D.-Y. Adjavon, M. Cook, M. Gkantia, K. J. Hayworth, G. B. Huang, W. T. Katz, F. Kämpf, Z. Lu, C. Ordish, T. Paterson, T. Stürner, E. T. Trautman, C. R. Whittle, L. E. Burnett, J. Hoeller, F. Li, F. Loesche, B. J. Morris, T. Pietzsch, M. W. Pleijzier, V. Silva, Y. Yin, I. Ali, G. Badalamente, A. S. Bates, R. J. Beresford, J. Bogovic, P. Brooks, S. Cachero, B. S. Canino, B. Chaisrisawat-suk, J. Clements, A. Crowe, I. d. H. Vicente, G. Dempsey, E. Donà, M. d. Santos, M. Dreher, C. R. Dunne, K. Eichler, S. Finley-May, M. A. Flynn, I. Hameed, G. P. Hopkins, P. M. Hubbard, L. Kiassat, J. Kovalyak, S. A. Lauchie, M. Leonard, A. Lohff, K. D. Longden, C. A. Maldonado, I. Moitra, S. S. Moon, C. Mooney, E. J. Munnelly, N. Okeoma, D. J. Olbris, A. Pai, B. Patel, E. M. Phillips, S. M. Plaza, A. Richards, J. R. Salinas, R. J. Roberts, E. M. Rogers, A. L. Scott, L. A. Scuderi, P. Seenivasan, L. S. Capdevila, C. Smith, R. Svirskas, S. Takemura, I. Tastekin, A. Thomson, L. Umayam, J. J. Walsh, H. Whittome, C. S. Xu, E. A. Yakal, T. Yang, A. Zhao, R. George, V. Jain, V. Jayaraman, W. Korff, G. W. Meissner, S. Romani, J. Funke, C. Knecht, S. Saalfeld, L. K. Scheffer, S. Waddell, G. M. Card, C. Ribeiro, M. B. Reiser, H. F. Hess, G. M. Rubin, and G. S. Jefferis, “Sexual dimorphism in the complete connectome of the drosophila male central nervous system,” bioRxiv, 2025.

[9] V. Lobato-Rios, S. T. Ramalingasetty, P. G. Özdil, J. Arreguit, A. J. Ijspeert, and P. Ramdya, “NeuroMechFly, a neuromechanical model of adult Drosophila melanogaster,” Nature Methods, vol. 19, pp. 620–627, May 2022.

[10] E. Todorov, T. Erez, and Y. Tassa, “Mujoco: A physics engine for model-based control,” in 2012 IEEE/RSJ International Conference on Intelligent Robots and Systems, pp. 5026–5033, IEEE, 2012.

[11] B. R. Cowley, A. J. Calhoun, N. Rangarajan, E. Ireland, M. H. Turner, J. W. Pillow, and M. Murthy, “Mapping model units to visual neurons reveals population code for social behaviour,” Nature, vol. 629, pp. 1100–1108, May 2024.

[12] J. K. Lappalainen, F. D. Tschopp, S. Prakhya, M. McGill, A. Nern, K. Shinomiya, S.-y. Takemura, E. Gruntman, J. H. Macke, and S. C. Turaga, “Connectome-constrained networks predict neural activity across the fly visual system,” Nature, vol. 634, pp. 1132–1140, Oct. 2024.

[13] S. Duan, L. L. Dong, and I. Fiete, “From synapses to dynamics: Obtaining function from structure in a connectome constrained model of the head direction circuit,” bioRxiv, pp. 2025–05, 2025.

[14] S. M. Pugliese, G. M. Chou, E. T. Abe, D. Turcu, J. K. Lancaster, J. C. Tuthill, and B. W. Brunton, “Connectome simulations identify a central pattern generator circuit for fly walking,” bioRxiv, pp. 2025–09, 2025.

[15] B. Szigeti, P. Gleeson, M. Vella, S. Khayrulin, A. Palyanov, J. Hokanson, M. Currie, M. Cantarelli, G. Idili, and S. Larson, “OpenWorm: an open-science approach to modeling Caenorhabditis elegans,” Frontiers in computational neuroscience, vol. 8, p. 137, 2014.

[16] S. J. Cook, T. A. Jarrell, C. A. Brittin, Y. Wang, A. E. Bloniarz, M. A. Yakovlev, K. C. Nguyen, L. T. Tang, E. A. Bayer, J. S. Duerr, et al., “Whole-animal connectomes of both Caenorhabditis elegans sexes,” Nature, vol. 571, no. 7763, pp. 63–71, 2019.

[17] P. Karashchuk, K. L. Rupp, E. S. Dickinson, S. Walling-Bell, E. Sanders, E. Azim, B. W. Brunton, and J. C. Tuthill, “Anipose: A toolkit for robust markerless 3d pose estimation,” Cell Reports, 2021.

[18] B. G. Pratt, S.-Y. J. Lee, G. M. Chou, and J. C. Tuthill, “Miniature linear and split-belt treadmills reveal mechanisms of adaptive motor control in walking Drosophila,” Current Biology, vol. 34, pp. 4368–4381.e5, Oct. 2024.

[19] C. Y. Zhang, Y. Yang, A. Sirbu, E. T. T. Abe, E. Wärnberg, E. J. Leonardis, D. E. Aldarondo, A. Lee, A. Prasad, J. Foat, K. Bian, J. Park, R. Bhatt, H. Saunders, A. Nagamori, A. R. Thanawalla, K. W. Huang, F. Plum, H. K. Beck, S. W. Flavell, D. Labonte, B. A. Richards, W. Brunton, E. Azim, B. P. Ölveczky, and T. D. Pereira, “MIMIC-MJX: Neuromechanical emulation of animal behavior,” 2025.

[20] H. Jaeger, “The “echo state” approach to analysing and training recurrent neural networks-with an erratum note,” Bonn, Germany: German national research center for information technology gmd technical report, vol. 148, no. 34, p. 13, 2001.

